# EFNB1 level affects drug response in DLBCL cell lines

**DOI:** 10.1101/2023.01.03.522579

**Authors:** Xiaoxi Li, Chenxiao Zhang, Minyao Deng, Zhengjin He, Hui Qian

## Abstract

Targeted therapy was a promising therapy for aggressive B-cell lymphoma. However, drug resistance was still an unavoidable problem. Here, we explored the effect of EFNB1 on drug response. Analysis of IC50 data showed that high level of EFNB1 was associated with resistance to most targeted drugs targeting BCR in human DLBCL cell lines. Drug response assay showed that Efnb1 could enhance SRC phosphorylation and increased the sensitivity of cells to SRC inhibitors. Meanwhile, Efnb1 could sensitize cells to most cytotoxic drugs, especially DOX and VCR. Efnb1 could confer cells sensitivity to VCR by enhancing the phosphorylation of Stmn1 at serine 28. Survival analysis revealed that the expression level of genes phosphorylated in Efnb1 cells were associated with poor prognosis of human DLBC. Together, this study revealed that EFNB1 level, as a non-genetic mechanism, greatly affected drug response of targeted drugs and cytotoxic drugs through the phosphorylation network. Our findings would provide important insights into EFNB1-SRC activated B-cell lymphoma and efficacy evaluation.

## Introduction

Lymphoma was a highly heterogeneous B-lymphoid hyperplasia disease. Although R-CHOP regimen had achieved good therapeutic effect in diffused large B-cell lymphoma (DLBCL), about 40% of patients still could not benefit. Extranodal dissemination was a typical feature of aggressive B-cell lymphoma, such as DLBCL(1), Burkitt’s lymphoma (BL)(2) and high-grade B-cell lymphoma (HGBL)(3,4), and it was also the main cause of treatment failure. The latest LymphGen(5) classification further divided DLBCL into 7 subtypes based on the hallmarks of genetic variations. The genetic hallmarks of MCD subtype included MYD88^L265P^, CD79B mutations, leading to activation of B-cell receptor (BCR) signaling pathway, and deletion of tumor suppressor gene CDKN2A. Extranodal lymphoma mainly belonged to the MCD subtype(5,6).

Targeted inhibition of BCR signaling pathway was the major development direction of targeted drugs. Ibrutinib, a Bruton’s tyrosine kinase (BTK) inhibitor, targeting BCR signaling, had substantial activity in the treatment of primary CNS lymphoma (PCNSL) (7,8). Ibrutinib plus R-CHOP also had been observed to enhance the survival benefit in MCD DLBLC(9). However, drug resistance had been observed in multiple mechanisms(10). Especially, resistance to BTK inhibitors could be caused by activation of pro-survival pathway, such as PI3K-AKT and others. Hence, identification of novel genes in extranodal lymphoma and its drug response profiles would be of great significance for understanding the pathogenesis of lymphoma and overcoming drug resistance.

Our previous study revealed that Efnb1 could promote extranodal dissemination of lymphoma in *Eμ-Myc;Utx^-/-^* mouse and high expression of Efnb1 was significantly associated with poor prognosis of human DLBCL(11). EFNB1 (Ephrin B1) was a single transmembrane tyrosine kinase and a ligand of Eph-Ephrin family. The multimers or oligomers of ephrin were required to form the Eph-Ephrin clusters and activation of Eph-Ephrin signaling(12). Moreover, the different size and conformation of Eph-Ephrin cluster had different quality of cellular responses(13,14). Hence, different expression level of ephrin may affect the structure and the strengthen of Eph-Ephrin signaling, and thus had different effect on drug responses.

Considering the tyrosine kinase activity of EFNB1, we guessed that EFNB1 had a crosstalk with BCR or other pro-survival pathways and determined the efficacy of targeted drugs. In this study, we explored the effect of EFNB1 on drug response of targeted drugs and chemotherapy. Our study found that the expression level of EFNB1 was closely correlated with drug response in DLBLC cell lines. Based on these findings, we preliminarily proposed individualized treatment strategy for DLBCL.

## Materials and methods

### Data acquisition and correspondence analysis

Drug IC50 dataset of drug panel and expression dataset of gene panel of DLBC cell lines were download from Genomics of Drug Sensitivity in Cancer (GDSC) (15) and Cancer Cell Line Encyclopedia (CCLE) (16). Correspondence analysis of drug IC50 data and gene expression data were performed using FactoMineR package in R Studio.

### Cell lines and constructs

*Eμ-myc;Cdkn2a^-/-^* murine B cell line was cultured in 45%DMEM, 45%IMDM, 10% fetal bovine serum, supplemented with 100 U/ml penicillin and streptomycin, 25μM β-mercaptoethanol. *Eμmyc;Cdkn2a^-/-^* murine B cell line was a kind gift from Prof. Hai Jiang at Center for Excellence in Molecular Cell Science. PCR-based test for mycoplasma contamination was performed every two weeks.

Gene of interest (GOI) with Kozak sequence GCCACC were amplified from cDNA of MA cells and cloned into retroviral vector MLS (LTR-MCS-SV40-GFP). Efnb1 was cloned into MSCV-based retroviral vectors MLS as previously(11). Ectopic expression constructs of Stmn1, Stmn1 S28A were established in this study. MSCV retroviral vectors with helper plasmid were co-transfected into HEK293T cells to produce retrovirus. GOI-GFP stable cell lines were established by retrovirus infection with polybrene(20ug/ml). For gradient dose analysis, fluorescence-activated cell sorting (FACS) was performed using FACSAria II (BD) to sort the GFP positive cells.

### Chemicals

Dasatinib (S1021), Ibrutinib (S2680), MK-2206 (S1078), Cytarabine (Ara-C, S1648), Methotrexate (MTX, S1210), Gemcitabine (GEM, S1714), Topotecan (TPT, S1231), Doxorubicin (DOX, S1208), Vincristine (VCR, S1241), Cisplatin (CDDP, S1166), Oxaliplatin (OXA, S1224), Decitabine (DAC, S1200) were purchased from Selleck.

All chemicals but CDDP and OXA were dissolved in dimethyl sulfoxide (DMSO) to concentrations of 10mM and aliquoted and stored at −20°C. CDDP and OXA were dissolved in Dimethylformamide (DMF) to concentrations of 10mM and aliquoted and stored at −80°C. Working concentrations for all chemicals were determined by lethal dose tests.

### Drug response analysis

GFP competition assay was performed as previous(17). Retrovirus infected cells with 30%-50% of GFP proportion were used in GFP competition assay. 4×104 cells in 100μl of BCM and 100μl 2x working solution diluted by BCM were mixed and incubated. 200μl and 400μl of fresh BCM were added at 24h and 48h. 100μl cell suspension were taken out to analysis the cell viability and GFP% of untreated and treated at 48h and 72h.

In GFP competition assay, cell viability was mainly used to evaluate whether the drug concentration achieved the effective lethal dose (LD). Drug concentration achieving LD80 to LD90 at 48h was optimum for most drugs in GFP competition assay. The living cells of PI negative population were gated to analysis GFP% of untreated and treated. GFP% of untreated and treated at 48h or 72h was used to calculated the Resistance Index (RI), which was used to evaluate the effect of genetic modification on therapeutic response of cells to tested drugs.

RI=(G1-G1*G2)/(G2-G1*G2). G1, GFP% in untreated. G2, GFP% in treated.

Gradient dose analysis was performed as follows. EV cells or sorted Efnb1 cells were mixed with 2x drug working solution and incubated as GFP competition assay. 200μl of fresh BCM were added at 24h. 100μl cell suspension were taken out to analysis the cell viability at 48h.

### Western blot analysis

Protein was extracted by RIPA buffer containing protease and phosphatase inhibitors. Protein samples were equally loaded with 25 μg protein on 10% gels and separated at 120V. PVDF membranes with transferred proteins were blocked with 5% milk in TBST and immunoblotted with the following antibodies: Phospho-Src (Tyr416) Rabbit mAb (#6943, CST, 1:1000), mouse monoclonal anti-ACTB (AC026, ABclonal, 1:100000). Chemiluminescent were detected by Amersham Imager 600 (GE Life Sciences).

### Mass spectrometry data analysis and pathway enrichment analysis

Cells were harvested by centrifugation and washed with cooled PBS three times. The precipitates were quick freezing at −80 and shipped to biotech company for mass spectrometry analysis with dry ice. Each sample was collected three times as three biological replicates.

The mass spectrometry analysis process was as follows. First, proteins were extracted and the concentrations of proteins were measured with BCA method. After trypsin enzymatic hydrolysis, the processed samples of replicates were combined and analyzed by LC-MS/MS to obtain raw files of the original mass spectrometry results. After MaxQuant (1.6.2.10) analysis, match the data and obtain the identification results.

Pathway enrichment analysis of DPs and DEPs were performed with Metascape(18). Pathway enrichment analysis has been carried out with GO Biological Processes. All genes in the genome have been used as the enrichment background.

### Prognostic analysis

Survival analysis was performed with the online tool SurvExpress(19). A human DLBCL dataset (Lenz Staudt Lymphoma GSE10846(20), n = 420) was chosen to survival analysis. The prognostic index (PI) was calculated by the expression value and the Cox model to generate the risk groups. The optimization algorithm was applied in risk grouping. The SurvExpress program was performed according to the tutorial.

### Statistical analysis

Depending on the type of experiment, unpaired t-test and log-rank test was used as indicated in figure legends. P values of < 0.05 were considered significant. *P < 0.05, **P < 0.01, ***P < 0.001, ****P < 0.0001.

## Results

### EFNB1 affects the response of targeted drugs targeting BCR signaling pathway

Abnormal activation of BCR signaling pathway was the feature of MCD DLBCL and targeted drugs were developed to block and kill lymphoma cells. The heterogeneity of lymphoma leaded to the difference on drug response. If lymphoma cells were sensitive to targeted drugs, it indicated that the survival of lymphoma cells were dependent on the pathway the drug targeted. To explore the correlation of EFNB1 level and activation of BCR signaling pathway, we analyzed IC50 data of kinase inhibitors targeting BCR pathway in DLBC cell lines.

On the whole, according to the expression level of EFNB1, cells could be divided into two types. Cells with low expression level of EFNB1 were sensitive to targeted drugs, while cells with high expression level of EFNB1 were resistant to targeted drugs. Specifically, according to the drug response profile and EFNB1 expression level, we divided the 24 cell lines into 5 types, A-E **(Figure 1A-B)**. EFNB1^low^ cell lines (TPM<=1.24), classified as Type D, were sensitive to most targeted drugs but ABL-BTK-SYK inhibitors, indicating that ABL-BTK-SYK pathway was not activated in the EFNB1^low^ cell lines. EFNB1^medium^ cell lines (1.37<=TPM<=1.88), classified as Type C, were resistant to all targeted drugs, indicating that the EFNB1^medium^ cell lines were not dependent on any one. Due to the difference on PI3K-AKT inhibitors, EFNB1^high^ cell lines (TPM>=1.94) were divided into two types, sensitive Type A and resistant Type B. In addition, the RL cell line had unique drug response pattern and was classified as Type E.

**FIGURE 1.**
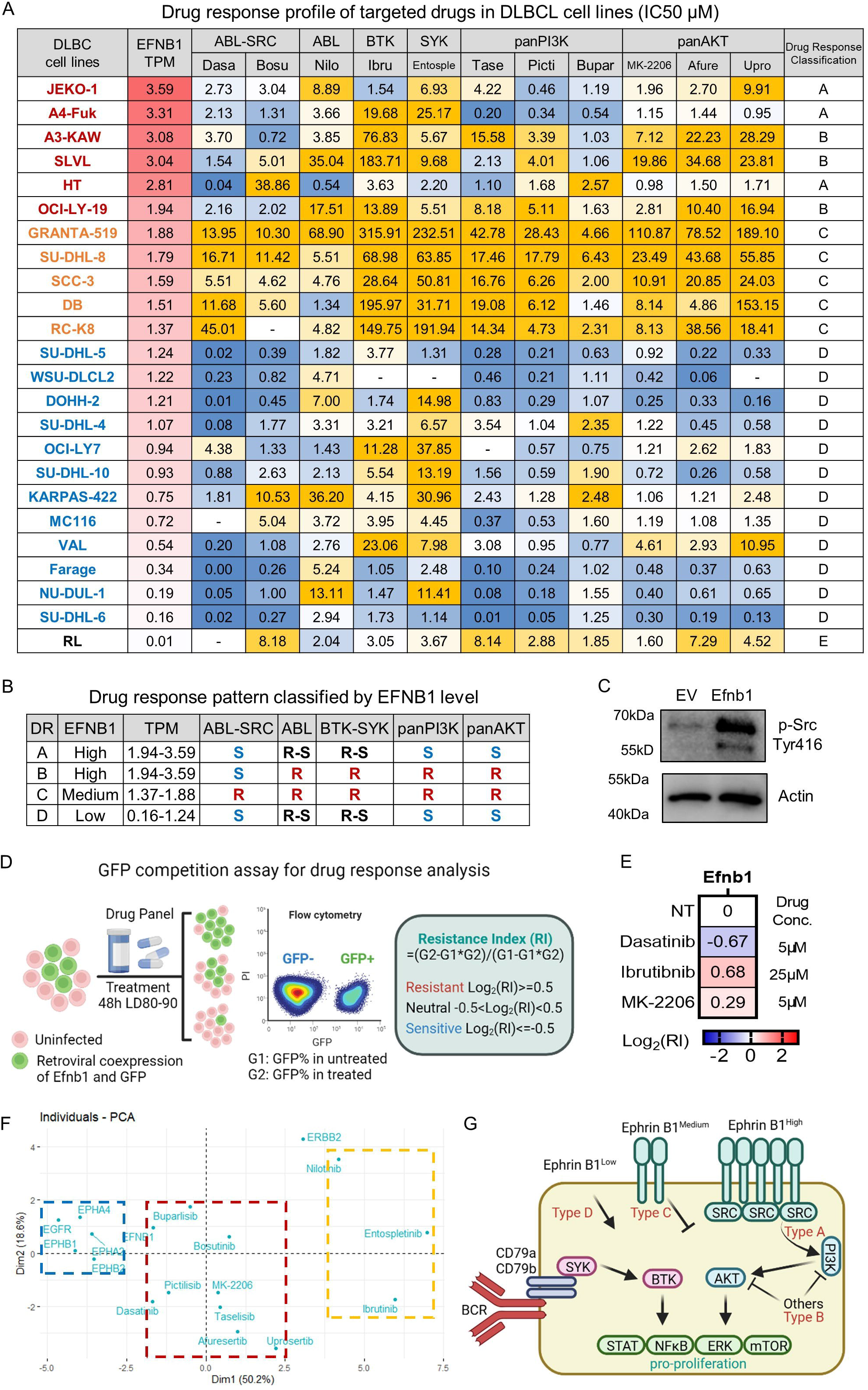
EFNB1 level is associated with the response of SRC-PI3K-AKT inhibitors in DLBC cell lines. **(A)**. Drug response profile of targeted drugs in DLBCL cell line. Efnb1 level was evaluated by TPM. TPM, Transcripts Per Kilobase per Million mapped reads. IC50, the half maximal inhibitory concentration. 24 DLBCL cell lines were divided into three groups: EFNB 1^high^, EFNB1^medium^ and EFNB1^low^. Corresponding to drug response pattern, the DLBCL cell lines were divided into four types, of which the EFNB1^high^ group was divided into type A and B, the EFNB1^medium^ group was type C, and the EFNB1^low^ group was type D. **(B)**. Drug response pattern of 4 types in DLBCL cell lines. **(C)**. Western bolt analysis of the phosphorylation level of SRC at tyrosine 416 site. Principal component analysis of EFNB1 expression and the response of kinase inhibitors in DLBCL cell lines. **(D)**. Diagram of GFP competition assay. Retroviral Efnb1 and GFP co-expressed cells were mixed with uninfected cells for drug treatment. The drug concentration was set according to the killing effect of drug treatment for 48h, usually reaching LD80-90. LD, Lethal dosage, evaluated by cell viability (PI-%). The GFP ratio of untreated cells and treated cells was analyzed at 72h. G1 represented the GFP ratio of untreated cells, and G2 represented the GFP ratio of drug treated cells. Calculate the Resistance Index (RI), RI=(G2-G1 * G2)/(G1-G1 * G2), and evaluate the effect of genes on drug response. Created with BioRender.com **(E)**. GFP competition assay of EFNB1 on targeted drugs. Log2 (RI)>=0.5 was considered as resistance, Log2 (RI)<=-0.5 was considered as sensitivity, otherwise it was neutral. NT, untreated. **(F)**. The principal component analysis was performed based on the expression level of EFNB1 (TPM) and IC50 of kinase inhibitors in DLBCL cell lines. Red dotted box, SRC-PI3K-AKT kinase inhibitors.Yellow dotted box, ABL-BTK-SYK kinase inhibitors. Blue dotted box, genes with similar expression patterns. **(G)**. Diagram of EFNB-SRC-PI3K-AKT axis. PI3K-AKT was activated through EFNB-SRC dependent manner in Type A cells or EFNB-SRC independent manner in Type D cells. Created with BioRender.com

Due to be sensitive to ABL-SRC inhibitor (Dasatinib) but ABL inhibitor (Nilotinib), we guessed that EFNB1^high^ cell lines were sensitive to SRC inhibition and SRC was activated in EFNB1^high^ cell lines. As SRC was an interacting protein of Efnb1(21) and could be phosphorylated by Ephrin B(22), we analyzed the phosphorylation level of SRC in Efnb1 overexpressed cells. The results showed that ectopic expression of Efnb1 significantly enhanced phosphorylation of SRC at tyrosine 416 **(Figure 1C)**, suggesting that activation of EFNB1-SRC contributed to the sensitivity of EFNB1^high^ cell lines to ABL-SRC inhibitor (Dasatinib) but ABL inhibitor (Nilotinib).

To verify the effect of EFNB1 on targeted drugs, we performed GFP competition assay in *Eμ-Myc;Cdkn2a^-/-^* cell lines **(Figure 1D)**. The results showed that Efnb1 conferred cells sensitive to Dasatinib and resistant to Ibrutinib **(Figure 1E)**, which was consistent with drug response pattern of EFNB1^high^ cell lines. The results indicated that activation of EFNB1-SRC increased the dependency of cell survival on SRC related pathway, thus increasing the sensitivity of SRC related inhibitors.

To identify the specificity of EFNB1 on drug response, we performed correspondence analysis of the expression level of EFNB1 receptors(23) (Ephb1, Ephb2, Epha2, Epha4) and other two single transmembrane tyrosine kinase receptors (EGFR, HER2) and the drug response of targeted drugs. Principal component analysis showed that the expression pattern of EFNB1 was the closest to the IC50 pattern of SRC-PI3K-AKT inhibitors, and had little correlation with the IC50 of ABL-BTK-SYK inhibitors **(Figure 1F)**. The expression pattern of other tyrosine kinase receptors had little relationship with the IC50 of targeted drugs. The results indicated that the expression level of EFNB1 was a specific biomarker to predict drug response of targeted drugs targeting BCR signaling pathway.

In conclusion, human DLBCL cell lines could be divided into four types according to EFNB1 level and the drug response patterns. Different expression level of Efnb1 could form different conformation of Eph-Ephrin cluster and different signaling network **(Figure 1G)**. The EFNB1^high^ cell lines with activated EFNB1-SRC were sensitive to SRC inhibitors, including Type A and Type B, which had different dependence on PI3K-AKT pathway. The EFNB 1 ^medium^ cell lines (type C) were generally resistant to targeted drugs, indicating that the EFNB 1 ^medium^ cell lines were not dependent on the BCR-PI3K-AKT pathways for survival. The EFNB1^low^ cell lines in with inactivated EFNB1-SRC (type D) were sensitive to targeted drugs, indicating that the EFNB1^low^ cell lines were strongly dependent on the BCR-PI3K-AKT pathways to promote cell proliferation and survival.

### EFNB1 confers chemo-susceptibility

To investigate the effect of EFNB1 on chemotherapy, we analyzed the drug response pattern of cytotoxic drugs in DLBC cell lines. Cytarabine (Ara-C) and Methotrexate (MTX) in R-HyperCVAD, Doxorubicin (DOX) and Vincristine (VCR) in R-CHOP regimen, Gemcitabine (GEM) in R-GCVP regimen, Etoposide (VP-16) in R-CEPP regimen, Cisplatin (CDDP) in R-ICE regimen, Oxaliplatin (OXA) in DHAX regimen, VNB in GemVNB regimen were analyzed.

The results showed the number of EFNB1-high cell lines (TPM>=1.24) sensitive to Ara-C, MTX, DOX, VP-16, CDDP was higher than that of EFNB1-low cell lines (TPM<=1.22) **(Figure 2A-B)**. GFP competition assay were performed to verify the effect of EFNB1 on cytotoxic drugs. The results showed that Efnb1 overexpressed cells were hypersensitive to cytotoxic drugs, especially DOX and VCR in R-CHOP regimen **(Figure 2C)**. The gradient dose assay further verified the gene-drug interaction of Efnb1-DOX **(Figure 2D**) and Efnb1-VCR **(Figure 2E)**. Decitabine (DAC) was tested as neutral control **(Figure 2F)**.

**FIGURE 2.**
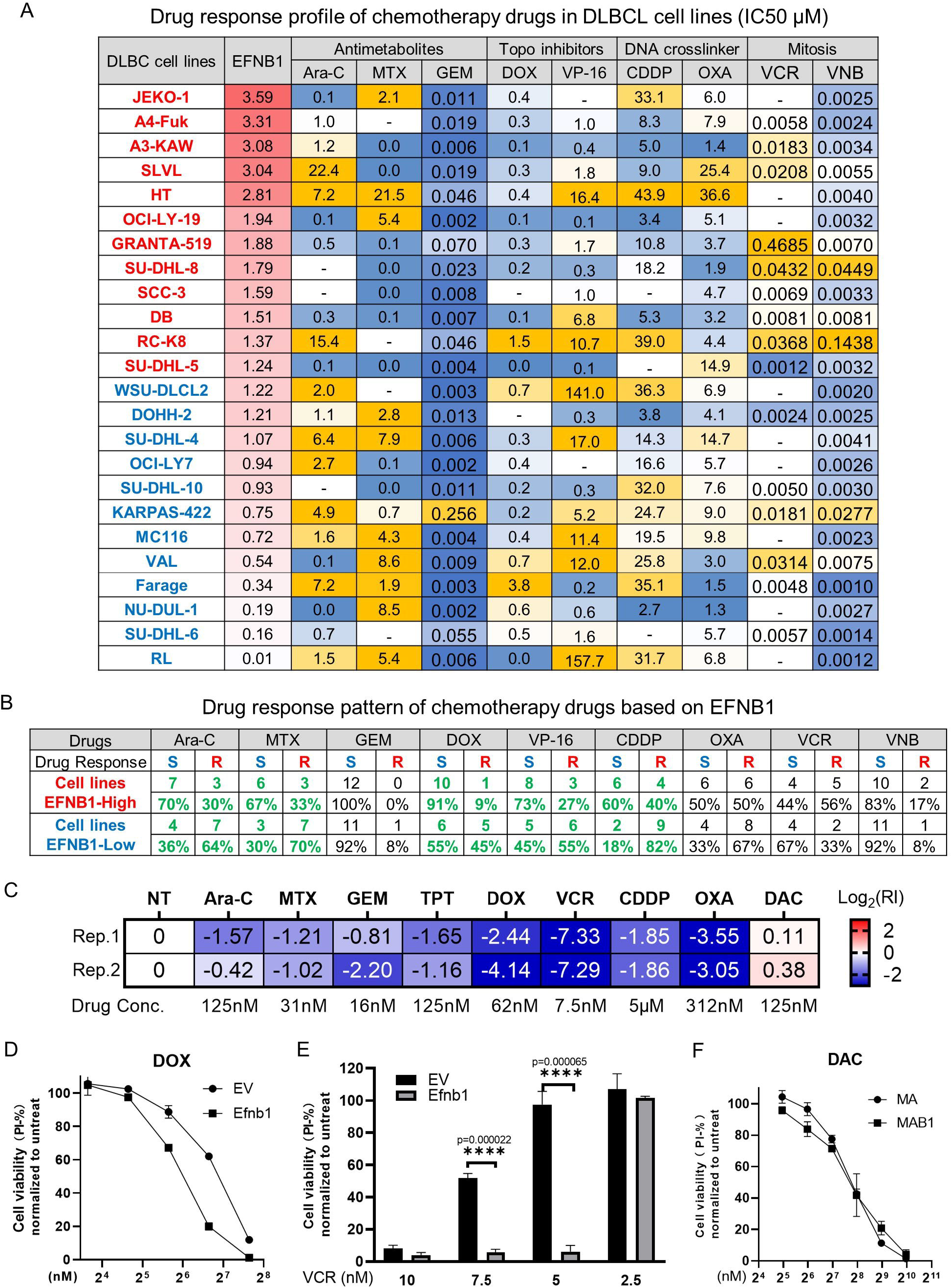
EFNB1 level is associated with the response of cytotoxic drugs in DLBC cell lines. **(A)**. Drug response pattern of chemotherapy drugs in DLBCL cell lines. The drug IC50 pattern of the DLBCL cell lines sorted by EFNB1 level. indicated lack of data. Yellow indicated tolerance and blue indicated sensitivity. **(B)**. The number and relative proportion of sensitive or drug-resistant cell lines in high and low EFNB1 cell lines. TPM of EFNB1 at 1.24-3.59 was high EFNB1 cell lines and TPM of EFNB1 less than 1.22 was low EFNB1 cell lines. The response pattern of the chemotherapy drugs shown in green was that the high EFNB1 cell lines tended to sensitivity and the low EFNB1 cell lines tended to resistance. **(C)**. GFP competition assay of EFNB1 on lymphoma drug panel. Log2 (RI)>=0.5 was considered as resistance, Log2 (RI)<=-0.5 was considered as sensitivity, otherwise it was neutral. NT, untreated. **(D)**. Gradient dose analysis of DOX on EV and Efnb1. Cell viability was measured by PI-% at 48 hours post of treatment. IC50_DOX_(EV)=140nM, IC50_DOX_(Efnb1)=68.6nM. **(E)**. Gradient dose analysis of VCR on EV and Efnb1. Cell viability was measured by PI-% at 48 hours post of treatment. unpaired t-test was used to test the significance. **(F)**. Gradient dose analysis of DAC on EV and Efnb1. Cell viability was measured by PI-% at 48 hours post of treatment.

Together, the results indicated that Efnb1 significantly enhanced sensitivity of cells to most cytotoxic drugs, suggesting that EFNB1 was a promising biomarker for responders to R-CHOP and other regimens.

### EFNB1 phosphorylation network involves in drug response and clinical prognosis

To understand the mechanism, we analyzed proteins and phosphorylated peptides in EV and Efnb1 cells by mass spectrometry. Totally, 35994 peptides involving 4135 proteins were identified in EV and Efnb1 cells. After filtered with the indicated parameters, 96 unique differential proteins (DPs), 91 differentially expressed proteins (DEPs), and 161 phosphorylated peptides were identified **(Figure 3A)**. Pathway enrichment analysis showed that EFNB1 significantly affected cell cycle and DNA repair signaling pathways **(Figure 3B)**.

**FIGURE 3.**
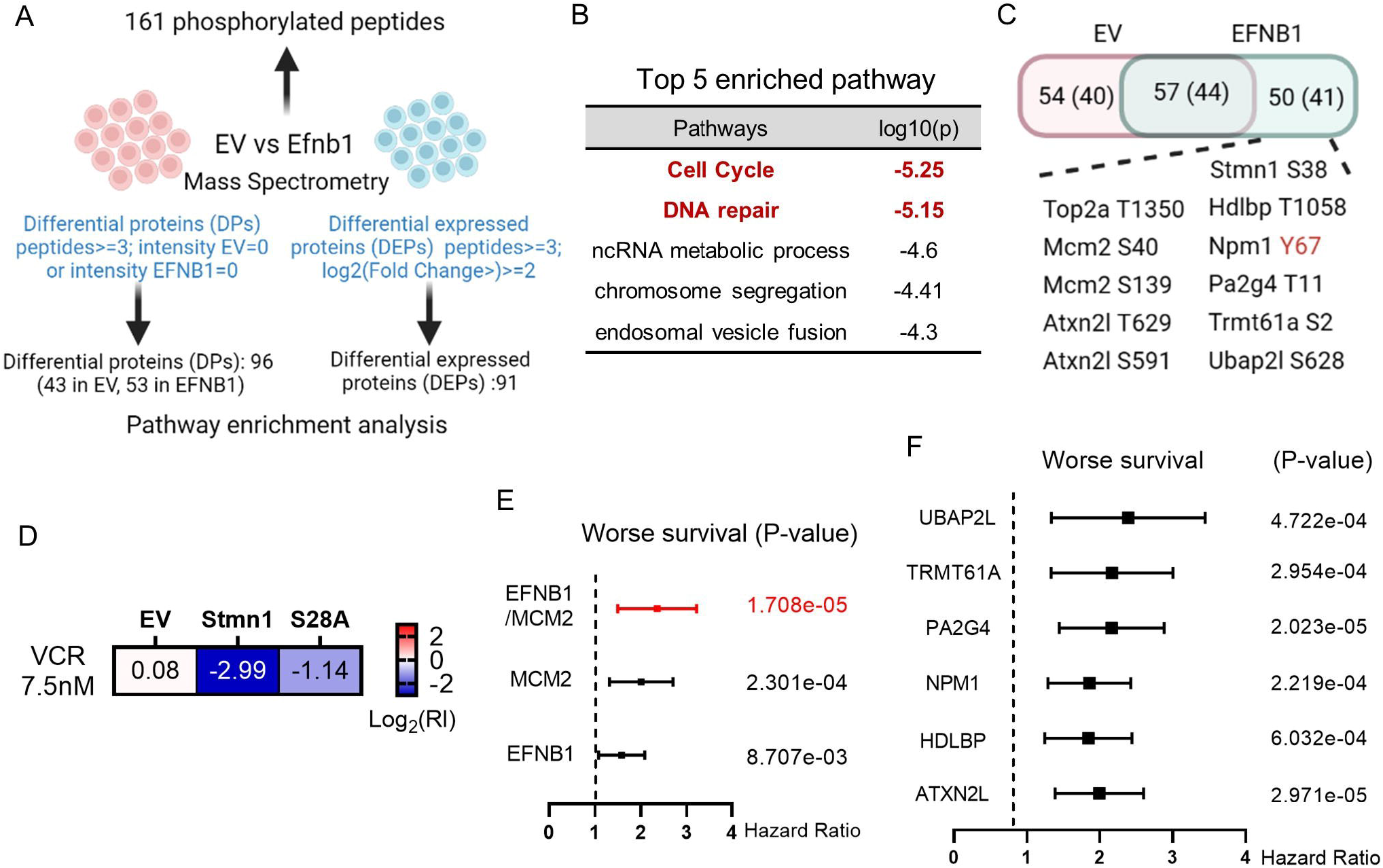
EFNB1 phosphorylated substrates involve in drug response and clinical prognosis. **(A)**. Mass spectrometry data analysis scheme of EV and Efnb11 cells. EV, empty vector. Differential proteins (DPs) referred to proteins only detected in EV or Efnb1, and the peptides>=3. Differently expressed proteins (DEPs) referred to proteins detected in both, and the peptides>=3, the relative intensity was more than 4-fold. 96 DPs, 91 DEPs, 161 phosphorylation sites/peptides (125 proteins) were obtained. Created with BioRender.com. **(B)**. Top 5 pathways enriched in DPs and DEPs. **(C)**. Distribution of 161 phosphorylation sites/peptides (125 proteins) in EV and Efnb1. Created with BioRender.com. **(D)**. GFP competition assay of wild-type Stmn1 and S28A mutant on VCR. **(E)**. Forest plot of the joint or individual expression of EFNB1 and MCM2 in 2 risk groups of human DLBCL. p-Value of the log-rank test were shown. **(F)**. Forest plot of phosphorylated substrates of EFNB1 in 2 risk groups of human DLBCL. p-Value of the log-rank test were shown. A human DLBCL dataset (Lenz Staudt Lymphoma GSE10846, n = 420) was chosen to survival analysis. The hazard ratio (HR), confidence interval, p-Value in forest plot were obtained from the SurvExpress program.

In EFNB1 cells, 50 phosphorylated peptides involving 41 proteins were identified **(Figure 3C)**, most of which involved drug response and poor prognosis. STMN1, a microtubule associated protein, was phosphorylation at Serine 28 in EFNB1 cells. Many reports showed the expression level of STMN1 was associated with chemoresistance of Vinca alkaloids and Taxanes. To investigate whether the phosphorylation of Serine 28 of STMN1 could affect the response of cells to VCR, we constructed the wild-type Stmn1 and S28A mutant cell lines to perform GFP competition assay. The results showed that ectopic expression of Stmn1 greatly sensitized cells to VCR, while the effect of S28A mutant on VCR was relatively weak than that of the wild-type **(Figure 3D)**. The results showed that increased phosphorylation of STMN1 at Serine 28 contributed to Efnb1 conferred sensitivity to VCR.

In addition, TOP2A was correlated to the sensitivity of topoisomerase inhibitors(24). MCM2-7, the helicase in DNA replication, were associated with poor prognosis of many cancers. Survival analysis showed that the joint expression of EFNB1 and MCM2 on the prognostic prediction was better than individual gene in human DLBCL **(Figure 3E)**. ATXN2L, HDLBP, NPM1, PA2G4, TRMT61A, UBAP2L were also significantly associated with poor prognosis of human DLBCL **(Figure 3F)**. The results indicated that EFNB1 phosphorylation network contributed to the progression of DLBCL.

Alteration of gene expression and protein modification, as non-genetic mechanisms, could regulate gene function, and thus greatly affected drug response and pathogenesis. Our finding suggested that EFNB1 could affect drug response and clinical prognosis through its expression level and the phosphorylation of its substrates.

### The clinical implication of EFNB1 on efficacy prediction and evaluation for DLBCL patients

Together, these evidences indicated that EFNB1 was a promising biomarker for efficacy evaluation in DLBCL. EFNB1^low^ cell lines were more sensitive to SRC-PI3K-AKT inhibitors and resistant to chemotherapy. EFNB1^Medium^ cell lines were resistant to most targeted drugs and sensitive to chemotherapy. EFNB1^High^ cell lines were divided into type A and type B. Type A was sensitive to PI3K-AKT inhibitors, and type B was resistant to PI3K-AKT inhibitors. Both were sensitive to SRC inhibitors. EFNB1^High^ cell lines had no obvious regularity in response to chemotherapy **(Figure 4A)**.

**FIGURE 4.**
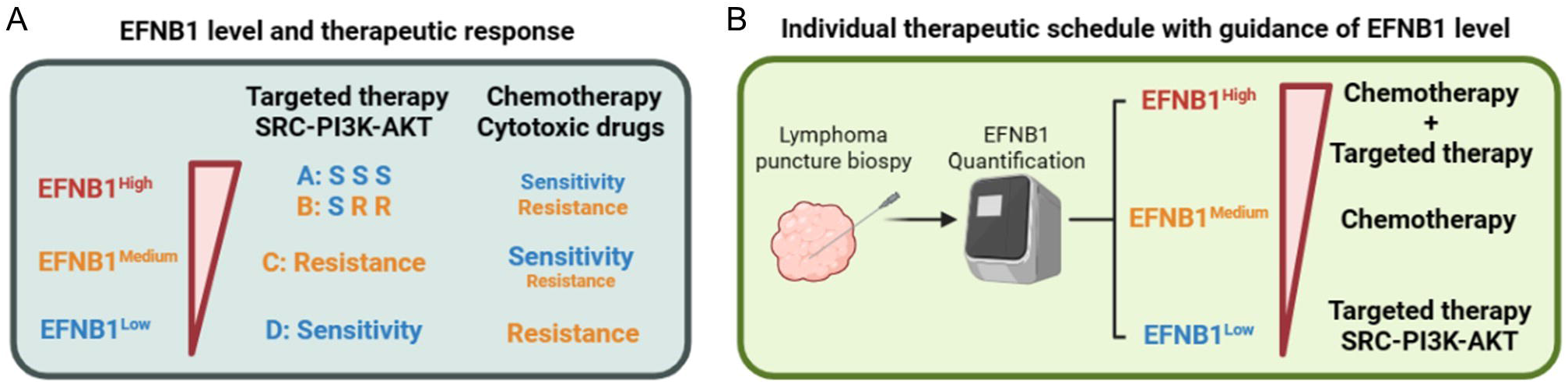
Proposal of EFNB1 guiding efficacy evaluation in DLBCL. (A). Therapeutic response of targeted drugs and chemotherapy drugs based on EFNB1 level. (B). Guiding principles of individualized treatment for DLBCL patients. Created with BioRender.com.

Therefore, we proposed a potential treatment strategy for DLBCL patients based on EFNB1 level to guide individualized rational therapy **(Figure 4B)**. We encouraged to evaluate the EFNB1 level of lymphoma biopsy from DLBLC patients before treatment. For patients with EFNB1^low^, targeted therapy targeting SRC-PI3K-AKT was suggested. For patients with EFNB1^medium^, chemotherapy was suggested. For patients with EFNB1^high^, due to the lack of biomarkers to distinguish type A and type B, targeted therapy combined with chemotherapy was suggested. Because lymphoma had been evolving, the evaluation of Efnb1 level were particularly available to patients with relapsed and/or refractory aggressive B-cell lymphoma.

## Discussion

BTK and PI3K were the main drug targets for BCR activated lymphoma. In this study, we found that the expression level of EFNB1 was correlated with the response profiles of BCR inhibitors, and EFNB1 could promote SRC phosphorylation. The characteristics of drug response profiles showed that EFNB1^low^ cells were sensitive to most BCR inhibitors, indicating that moderate and high expression of EFNB1 would inhibit BCR signals and weaken the dependence on BCR signals. Meanwhile, the drug response profiles revealed that inhibition of SRC could be a promising way to overcome drug resistance in EFNB1^high^ cells. Given that some of EFNB1^high^ cells were sensitive to PI3K-AKT inhibitors, biomarkers and mechanism of the responders to PI3K-AKT should be further identified to guide precise targeted drug therapy. The dosage effects of EFNB1 level on drug response could attribute to the diversity of Eph-Ephrin cluster(13,14).

BTK inhibitors had shown good effects in preclinical models and clinical trials, and could significantly improve the prognosis. In this study, we found that the lethal concentration of Ibrutinib on most DLBCL cell lines and murine cell line was too high, indicating that these cell lines were not dependent on BTK. On the one hand, this phenomenon might attribute to the lack of conditions for activation of BTK signaling pathway in culture. On the other hand, inhibition of cell proliferation might not be the main role of BTK inhibitors. Inhibiting cell adhesion and migration, and reducing extranodal dissemination of lymphoma should be the main reasons for BTK inhibitors to improve prognosis. Further *in vivo* therapeutic experiments were needed to verify the effect of EFNB1 on drug response.

Ectopic expression of EFNB1 could significantly enhance the sensitivity of cells to cytotoxic drugs, indicating that EFNB1 weakened the DNA repair pathways and enhanced cell death pathways. Mass spectrometry identified that several phosphorylation levels of proteins involved in DNA replication and DNA repair were highly expressed in EFNB1 cells, such as TOP2A, indicating that EFNB1 could affect chemotherapy sensitivity by regulating phosphorylation. However, the phosphorylation substrates of EFNB1 identified in our data were mainly the serine and the threonine, and only the phosphorylation site on NPM1 was the tyrosine. Given that EFNB1 greatly affected drug response, EFNB1 phosphorylation network and the crosstalk of EFNB1 and BCR signaling pathway should become the fucus of aggressive B-cell lymphoma in the future. The law between EFNB1 and the drug response should be further explored and proven by clinical evidence, which would significantly contribute to optimizing efficacy prediction and treatment of DLBCL patients.

The abnormal activation of BCR signaling pathway in MCD DLBCL was the basis for targeted therapy. However, the heterogeneity of lymphoma was high and dynamic, and drug resistance was still the major reason of treatment failure in clinical. Our findings highlighted the role of EFNB1 in evaluation of therapeutic response, especially targeted therapy. Our study would provide meaningful insights into the development and rational evaluation of targeted drugs in aggressive B-cell lymphoma.

## Data Availability Statement

All data and materials are presented in the main manuscript or supplementary materials and are available on request. The mass spectrometry proteomics data have been deposited to the ProteomeXchange Consortium via the PRIDE partner repository with the dataset identifier PXD038838.

## Author Contributions

XXL conceptualized, designed the experiments and wrote the manuscript. HQ helped with designing the experiments and reviewing the manuscript. CXZ performed the drug response assay. XXL and MYD analyzed the IC50 data of DLBC cell lines and the MS data. ZJH helped with the retrovirus package and established the stable cell lines. All authors discussed the results and commented on the manuscript. XXL and HQ jointly supervised the study.

